# Rarefaction is currently the best approach to control for uneven sequencing effort in amplicon sequence analyses

**DOI:** 10.1101/2023.06.23.546313

**Authors:** Patrick D. Schloss

## Abstract

Considering it is common to find as much as 100-fold variation in the number of 16S rRNA gene sequences found across samples in a study, researchers need to control for the effect of uneven sequencing effort. How to do this has become a contentious question. Some have argued that rarefying or rarefaction is “inadmissible” because it omits valid data. A number of alternative approaches have been developed to normalize and rescale the data that purport to be invariant to the number of observations. I generated community distributions based on 12 published datasets where I was able to assess the ability of multiple methods to control for uneven sequencing effort. Rarefaction was the only method that could control for variation in uneven sequencing effort when measuring commonly used alpha and beta diversity metrics. Next, I compared the false detection rate and power to detect true differences between simulated communities with a known effect size using various alpha and beta diversity metrics. Although all methods of controlling for uneven sequencing effort had an acceptable false detection rate when samples were randomly assigned to two treatment groups, rarefaction was consistently able to control for differences in sequencing effort when sequencing depth was confounded with treatment group. Finally, the statistical power to detect differences in alpha and beta diversity metrics was consistently the highest when using rarefaction. These simulations underscore the importance of using rarefaction to normalize the number of sequences across samples in amplicon sequencing analyses.

**Importance:** Sequencing 16S rRNA gene fragments has become a fundamental tool for understanding the diversity of microbial communities and the factors that affect their diversity. Due to technical challenges, it is common to observe wide variation in the number of sequences that are collected from different samples within the same study. However, the diversity metrics used by microbial ecologists are sensitive to differences in sequencing effort. Therefore, tools are needed to control for the uneven levels of sequencing. This simulation-based analysis shows that despite a longstanding controversy, rarefaction is the most robust approach to control for uneven sequencing effort. The controversy started because of confusion over the definition of rarefaction and violation of assumptions that are made by methods that have been borrowed from other fields. Microbial ecologists should use rarefaction.

## Introduction

The ability to generate millions of 16S rRNA gene amplicon and metagenomic sequence reads has allowed researchers to multiplex multiple samples on the same sequencing run by pooling separate PCRs that can be deconvoluted later based on index (aka barcode) sequences that are embedded into the sequence of the PCR primers (1–3). Unfortunately, it is common to observe variation in the number of sequence reads from each sample vary by as much as 100-fold (e.g, see Figure S1). This occurs because pooling of DNA from multiple PCRs is fraught with numerous opportunities for technical errors to compound leading to a skewed distribution. Aside from developing better methods of pooling DNA, the question of how to control for uneven sequencing effort in microbial ecology studies has become controversial.

The practice of rarefaction has been commonly used in ecology for more than 50 years as a tool to control for uneven sequencing effort across experimental replicates (4, 5). Microbial ecologists have used it to compare 16S rRNA gene sequence data for the past 25 years (6–8). With rarefaction the investigator selects a desired threshold of sequencing effort and removes any samples below that threshold. They then randomly select that many sequences, with replacement from each sample. Based on the observed sequence counts, the researcher can then calculate alpha diversity metrics including richness and diversity indices or beta diversity metrics such as a Jaccard or Bray-Curtis dissimilarity index. I refer to this single sampling as a subsample; this method is implemented as the `sub.sample` function in mothur (9) and the `rrarefy` function in the `vegan` R package (10). Rarefaction repeats the subsampling a large number of times (e.g., 100 or 1,000 times) and calculates the mean of the alpha or beta diversity metric over those subsamplings; rarefaction is implemented in mothur using the `summary.single` and `dist.shared` functions (9) and with the `vegan` R package using the `rarefy` or `avgdist` functions (10). Rarefaction effectively tells a researcher what an alpha or beta diversity metric would have been for a collection of samples if they were all sequenced to the same depth. Although a closed form equation exists to calculate the expected richness (5), it is computationally easier to empirically calculate richness and other alpha and beta diversity metrics by rarefaction.

In 2014, McMurdie and Holmes (11) announced that “rarefying” of microbial community data was “statistically inadmissible” because it omits valid data. In their simulations, they observed that rarefying reduced the statistical power to correctly cluster samples into the same treatment groups based on beta diversity metrics. Although the detail was lost on many subsequent researchers, they did not describe rarefaction, but rarefying (12). According to McMurdie and Holmes, rarefying is a single subsampling of an OTU abundance table whereas rarefaction repeats the subsampling step many times. Furthermore, their simulations penalized the rarefied data by removing 15% of the samples and evaluated clustering quality using a clustering algorithm that performed worse for rarefied data. According to my reanalysis of rarefaction using the original simulation framework, rarefaction outperformed other normalization method being used in 2014 for both alpha and beta diversity metrics (12). Others have also critiqued the original work (13). Regardless, there is a general sense in the community that rarefying and rarefaction should not be used with microbiome data.

Since McMurdie and Holmes published their simulations, others have developed alternative approaches to control for uneven sequencing effort in amplicon sequencing studies. For alpha diversity metrics, Willis used toy datasets to demonstrate that one could estimate the richness for each sample in a dataset and use those values for statistical comparisons (14). Non-parametric estimators of richness and diversity (15, 16) and parametric estimators of richness (17) have been used in microbial ecology studies. For beta diversity metrics at least four approaches have been pursued. First, one could use relative abundance values where the observed number of sequences in an OTU is divided by the total number of sequences in the sample (18). Second, normalization strategies have been developed where the relative abundance is multiplied by the size of the minimum desired sequencing effort and fractional values are reapportioned to the OTUs to obtain integer values (19, 20). Third, a variety of center log-ratio methods have been developed where the compositional nature of the OTU counts is removed and used to calculated Euclidean distances (aka Aitchison distances) (18, 20–23). This strategy is purported to control for uneven sequencing effort (22, 24); however, some have noticed that this feature breaks down under certain conditions (25). Finally, as mentioned above, variance stabilization transformations have been recommended to generate values that can be used to calculate Euclidean distances (11).

The ongoing controversy over the use of rarefaction and the recent development of alternative strategies to control for uneven sequencing effort caused me to question how these methods compared to each other using a simulation framework that overcame the issues with the McMurdie and Holmes study. My analysis included 16S rRNA gene sequence data from from 12 studies that characterized the variation in bacterial communities from diverse environments (Table 1 and Figure S1). The sequences were assigned to OTUs using a standard pipeline and their frequencies and the number of sequences found in each sample were used to generate simulated communities and treatment effects. For each dataset and simulation, 100 replicate datasets were generated and used as input to each of the strategies for controlling for uneven sequencing effort. My overall conclusion was that rarefaction outperformed the alternative strategies.

**Table 1.** Summary of studies used in the analysis. For all studies, when rarefaction was used the number of sequences used from each dataset was the size of the smallest sample. A graphical representation of the distribution of sample sizes for each dataset and the samples that were removed from each dataset are provided in Figure S1. This table is similar to Table 1 from (27).

| Dataset (Ref) | Samples | Total sequences | Median sample size | Mean sample size | Range of sample sizes | SRA study accession |
| --- | --- | --- | --- | --- | --- | --- |
| Bioethanol (35) | 95 | 3,970,972 | 16,014 | 41,799 | 3,690-356,027 | SRP055545 |
| Human (36) | 490 | 20,828,275 | 32,452 | 42,506 | 10,439-422,904 | SRP062005 |
| Lake (37) | 52 | 3,145,486 | 69,205 | 60,490 | 15,135-110,993 | SRP050963 |
| Marine (38) | 7 | 1,484,068 | 213,091 | 212,009 | 132,895-256,758 | SRP068101 |
| Mice (3) | 348 | 2,785,641 | 6,426 | 8,004 | 1,804-30,311 | SRP192323 |
| Peromyscus (39) | 111 | 1,545,288 | 12,393 | 13,921 | 4,454-33,502 | SRP044050 |
| Rainforest (40) | 69 | 936,666 | 11,464 | 13,574 | 4,880-37,403 | ERP023747 |
| Rice (41) | 490 | 22,623,937 | 43,399 | 46,171 | 2,777-192,200 | SRP044745 |
| Seagrass (42) | 286 | 4,135,440 | 13,538 | 14,459 | 1,830-45,076 | SRP092441 |
| Sediment (43) | 58 | 1,151,389 | 17,606 | 19,851 | 7,686-67,763 | SRP097192 |
| Soil (44) | 18 | 932,563 | 50,487 | 51,809 | 46,622-58,935 | ERP012016 |
| Stream (45) | 201 | 21,017,610 | 90,621 | 104,565 | 8,931-394,419 | SRP075852 |

## Results

### Without rarefaction, metrics of alpha diversity are sensitive to sequencing effort

To test the sensitivity of various approaches of measuring alpha diversity to sequencing effort, I generated null models using OTU counts and sequencing depths from 12 studies. Under a null model, each community from the same dataset would be expected to have the same alpha diversity regardless of the sequencing effort. I measured the richness of the communities in each dataset without any correction, using scaled ranked subsampling (SRS) normalized OTU counts, with estimates based on non-parametric and parametric approaches, and using rarefaction (e.g. Figure S2). For each dataset, all of the approaches, except for rarefaction, showed a strong correlation between richness and the number of sequences in the sample (Figure 1A). Next, I assessed diversity using the Shannon diversity index and the inverse Simpson diversity index without any correction, using normalized OTU counts, and rarefaction; I also used a non-parametric estimator of Shannon diversity. The correlation between sequencing depth and the diversity metric was not as strong as it was for richness and the inverse Simpson diversity values were less sensitive than the Shannon diversity values; however, the correlation to the diversity metrics calculated with rarefaction were the lowest for all of the metrics and studies (Figure 1A). The alpha-diversity metrics calculated with rarefaction consistently demonstrated a lack of sensitivity to sequencing depth.

**Figure 1.**
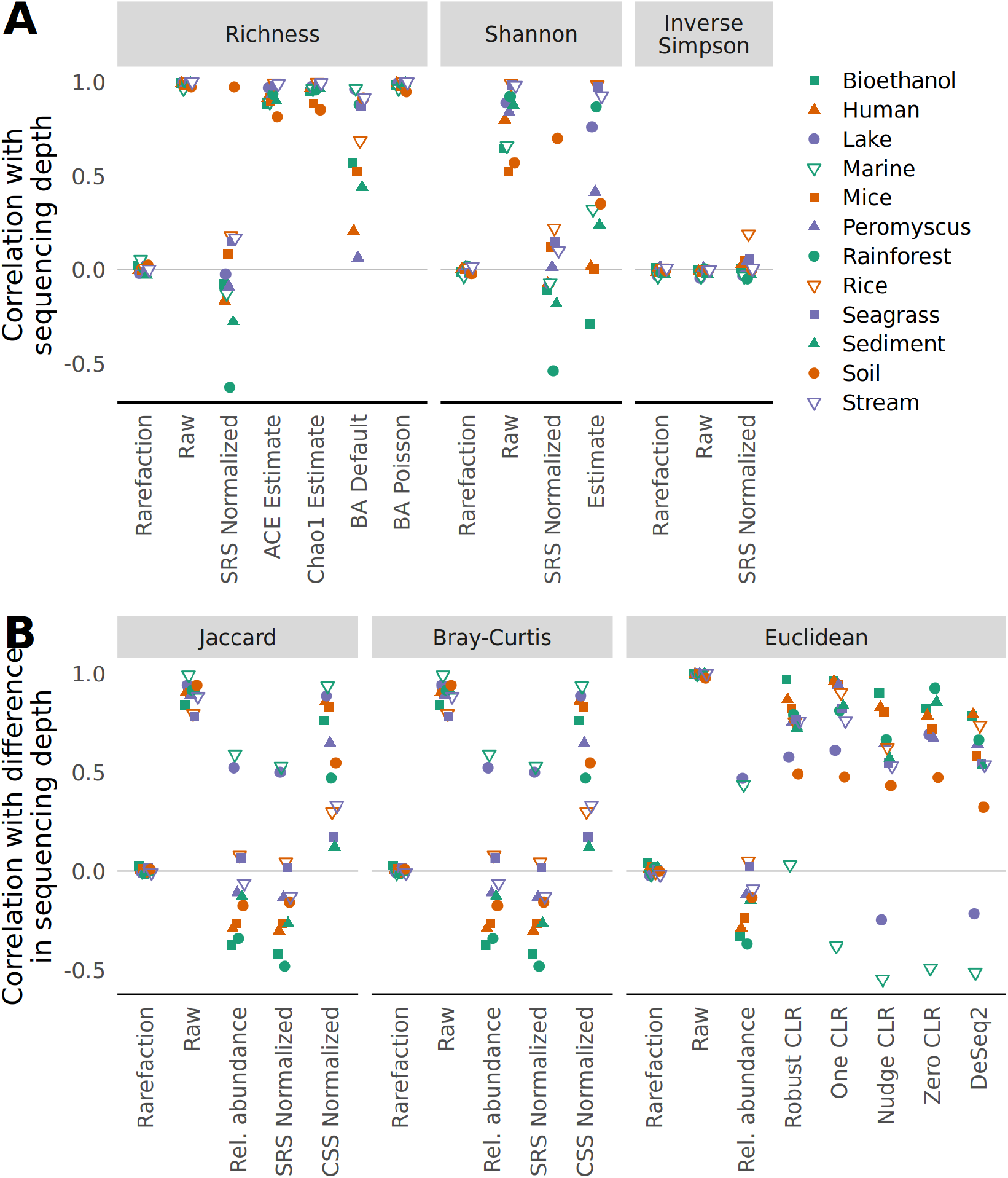
Rarefaction eliminates the correlation between sequencing depth and alpha diversity (A) and between differences in sequencing depth and beta (B) diversity metrics when using null community models. Examples of the relationship between different metrics and methods for controlling for uneven sequencing effort are provided in Figures S2 and S3 for alpha and beta diversity metrics, respectively. Each point represents the mean of 100 random null community models; the standard deviation was smaller than the size of the plotting symbol.

### Without rarefaction, metrics of beta diversity are sensitive to sequencing effort

To test the sensitivity of various approaches of measuring beta diversity to sequencing effort, I used the same null models used for studying the sensitivity of alpha diversity. Under a null model, the ecological distance between any pair of samples would be the same regardless of the difference in the number of sequences observed in each sample (e.g., Figure S3). First, I calculated the Jaccard distance coefficient between all pairs of communities within a dataset. The Jaccard distance coefficient is the fraction of OTUs that are unique to either community and does not account for the abundance of the OTUs. Jaccard distances were calculated using the uncorrected OTU counts, with rarefaction, relative abundances, and following normalization using cumulative sum scaling (CSS) and SRS. Only the distances calculated with rarefaction showed a lack of sensitivity to sequencing effort (Figure 1B). Second, I analyzed the sensitivity of the Bray-Curtis distance coefficient, which is a popular metric that incorporates the abundance of each OTU. Similar to what I observed with the Jaccard coefficient, only the data normalized with rarefaction showed a lack of sensitivity to sequencing effort (Figure 1B). Third, I calculated the Euclidean distance on raw OTU counts where the central log-ratio (CLR) was calculated (i.e., Aitchison distances) by ignoring OTUs in samples with zero counts (Robust CLR), adding a pseudo-count of one to all OTU counts prior to calculating the CLR (One CLR), adding a pseudo-count of one divided by the total number of sequences obtained for the community (Nudge CLR), and imputing the value of zero counts (Zero CLR). The Aitchison distances were all strongly sensitive to sequencing effort (Figure 1B). Finally, I used the variance-stabilization transformation (VST) from DESeq2 prior to calculating Euclidean distances. Again, there was a strong sensitivity to sequencing effort (Figure 1B). Although Euclidean distances are not typically used on raw or rarefaction-normalized count data in ecology (26), Euclidean distances calculated with rarefaction were not sensitive to sequencing effort. Across each of the beta diversity metrics and approaches used to account for uneven sequencing effort and sparsity, rarefaction was the least sensitive approach to differences in sequencing effort.

### Rarefaction limits the detection of false positives when sequencing effort and treatment group are confounded

Next, I investigated the impact of the various strategies and metrics on falsely detecting a significant difference using the the same communities generated from the null model in the analysis of alpha and beta diversity metrics. To test for differences in alpha and beta diversity I used the non-parametric Wilcoxon test and non-parametric permutation-based multivariate analysis of variance (PERMANOVA). First, I employed an unbiased null treatment model to measure the false detection rate, which should not have meaningfully differed from 5%. Indeed, for each dataset and alpha and beta diversity metric and strategy for accounting for uneven sequencing, approximately 5% of the tests yielded a significant result (Figure 2). Second, I employed a biased null treatment model where the treatment group was completely confounded with the number of sequences in each sample. Under this model, only the data normalized with rarefaction consistently resulted in a 5% false positive rate for alpha and beta diversity metrics (Figure 2). These results aligned with the observed sensitivity of alpha and beta diversity metrics to sequencing effort and underscore the value of rarefaction.

**Figure 2.**
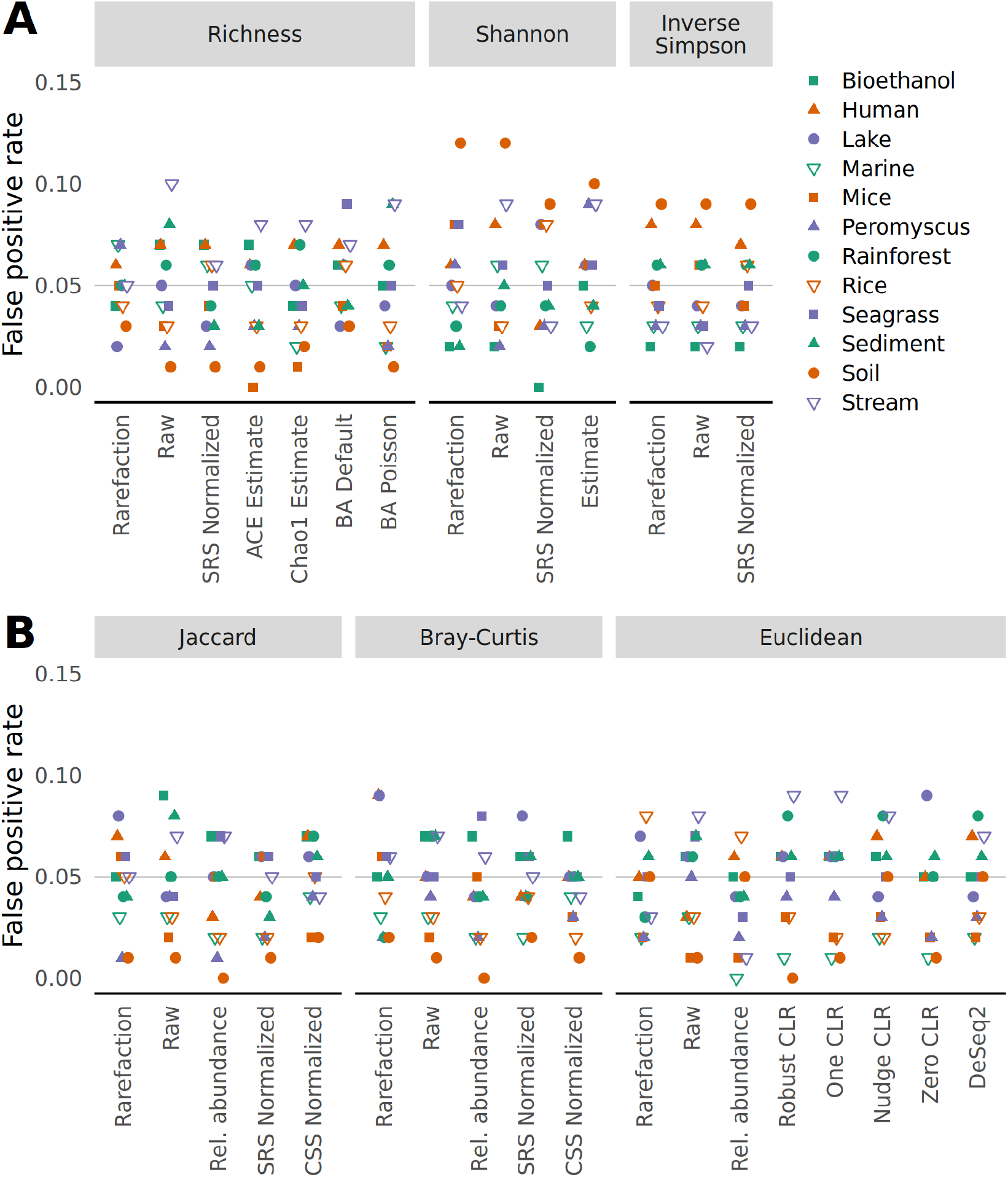
The risk of falsely detecting a difference between treatment groups drawn from a null model does not meaningfully vary from 5%, regardless of approach for controlling for uneven sequencing depth. Samples were randomly assigned to different treatment groups. To calculate the false detection rate, datasets were regenerated 100 times and differences in alpha diversity were tested using a Wilcoxon test (A) and differences in beta diversity were tested using PERMANOVA (B) at a 5% threshold. The false positive rate was the number of times a dataset yielded a significant result.

### Rarefaction preserves the statistical power to detect differences between treatment groups

To assess the impact of different approaches to control for uneven sequencing effort I performed two additional simulations. In the first simulation, I implemented a skewed abundance distribution model to create two treatment groups for each dataset that were each populated with half of the samples each with the same number of sequences as the samples had in the observed data. The two treatment groups varied in their structure such that one had the same abundances as the null distribution above and the other had 10% of its OTUs randomly selected to increase their counts by 5%. The power to detect differences in richness between the two simulated treatment groups by all approaches was low (Figure 4A). This was likely because the approach for generating the perturbed community did not necessarily change the number of OTUs in each treatment group. Regardless, the simulations testing differences in richness using the Rice and Stream datasets had the greatest power when the richness data were calculated with rarefaction. To explore this further, a richness-adjusted community model was created by removing 3% of the OTUs from a null model model. As suggested by the first simulation, the richness data calculated with rarefaction had a higher statistical power than the other approaches when measuring richness (Figure 5). The simulations testing the power to detect differences in Shannon diversity also showed that rarefaction performed better than the other methods (Figure 4A). When testing for differences in the Inverse Simpson diversity index the the difference between rarefaction and the other methods was negligible (Figure 4A). For tests of beta diversity I found that rarefaction was the most reliable approach to maintain statistical power to detect differences between two communities. Among the tests using the Jaccard and Bray-Curtis metrics, raw count data and CSS normalized data had little power relative to using rarefaction, relative abundance, and SRS to normalize the uneven sequencing depths. The differences in power between counts normalized with rarefaction, relative abundance, and SRS data was small, but if there were differences, the power obtained using rarefaction was greater than the other methods. Among the tests using Euclidean distances, using raw counts and CLR and DESeq2 transformed data had little power relative to the distances calculated using rarefaction and relative abundances. This power-based analysis of the simulated communities using different methods of handling uneven sample sizes demonstrated the value of rarefaction for preserving the statistical power to detect differences between treatment groups for measures of alpha and beta diversity.

**Figure 3.**
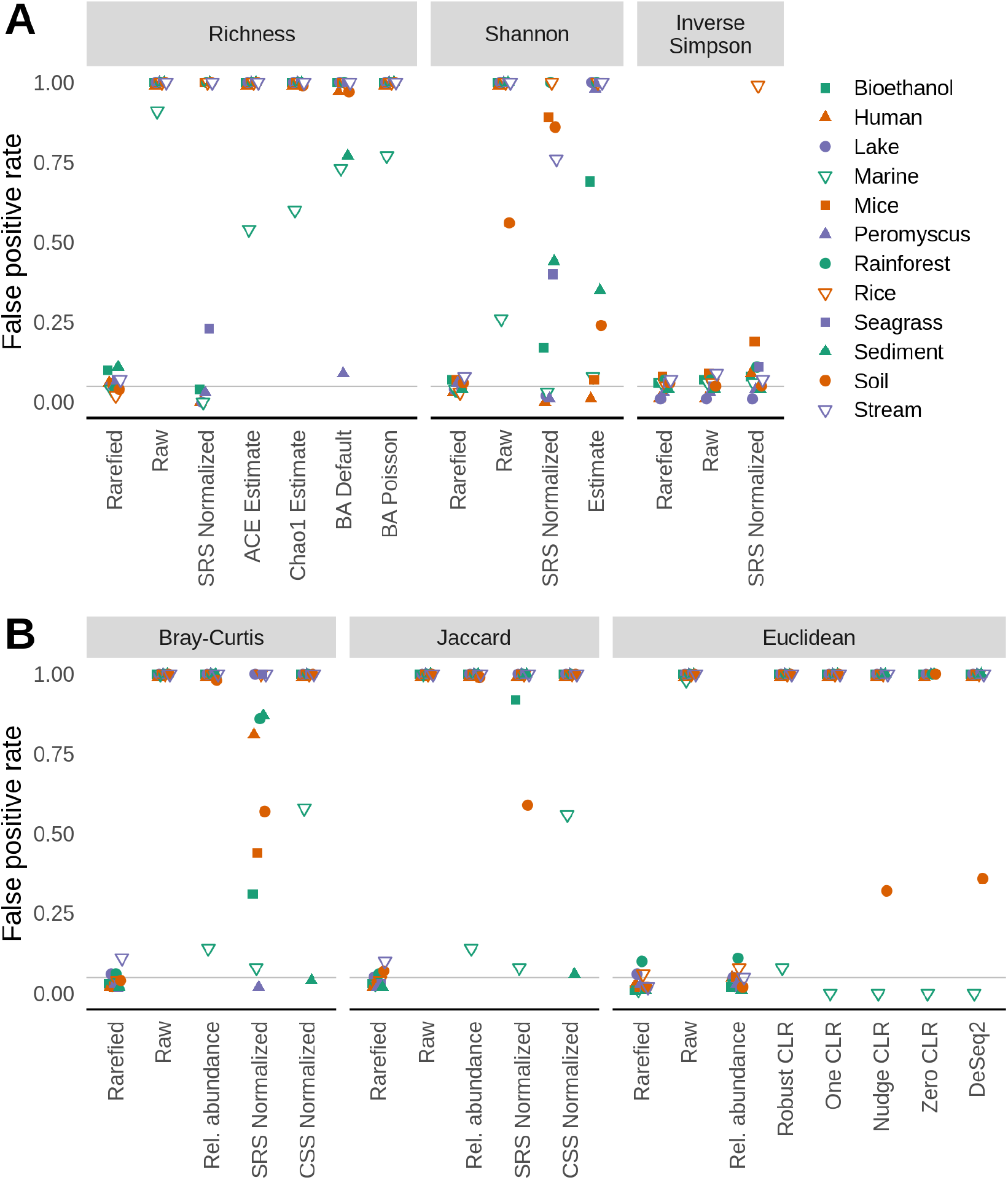
The risk of falsely detecting a difference between treatment groups drawn from a null model does not meaningfully vary from 5% when data are normalized by rarefaction when sequencing depth is confounded with treatment group. Samples were assigned to different treatment groups based on whether they were above the median number of sequences for each dataset. To calculate the false detection rate, datasets were regenerated 100 times and differences in alpha diversity were tested using a Wilcoxon test (A) and differences in beta diversity were tested using PERMANOVA (B) at a 5% threshold. The false positive rate was the number of times a dataset yielded a significant result.

**Figure 4.**
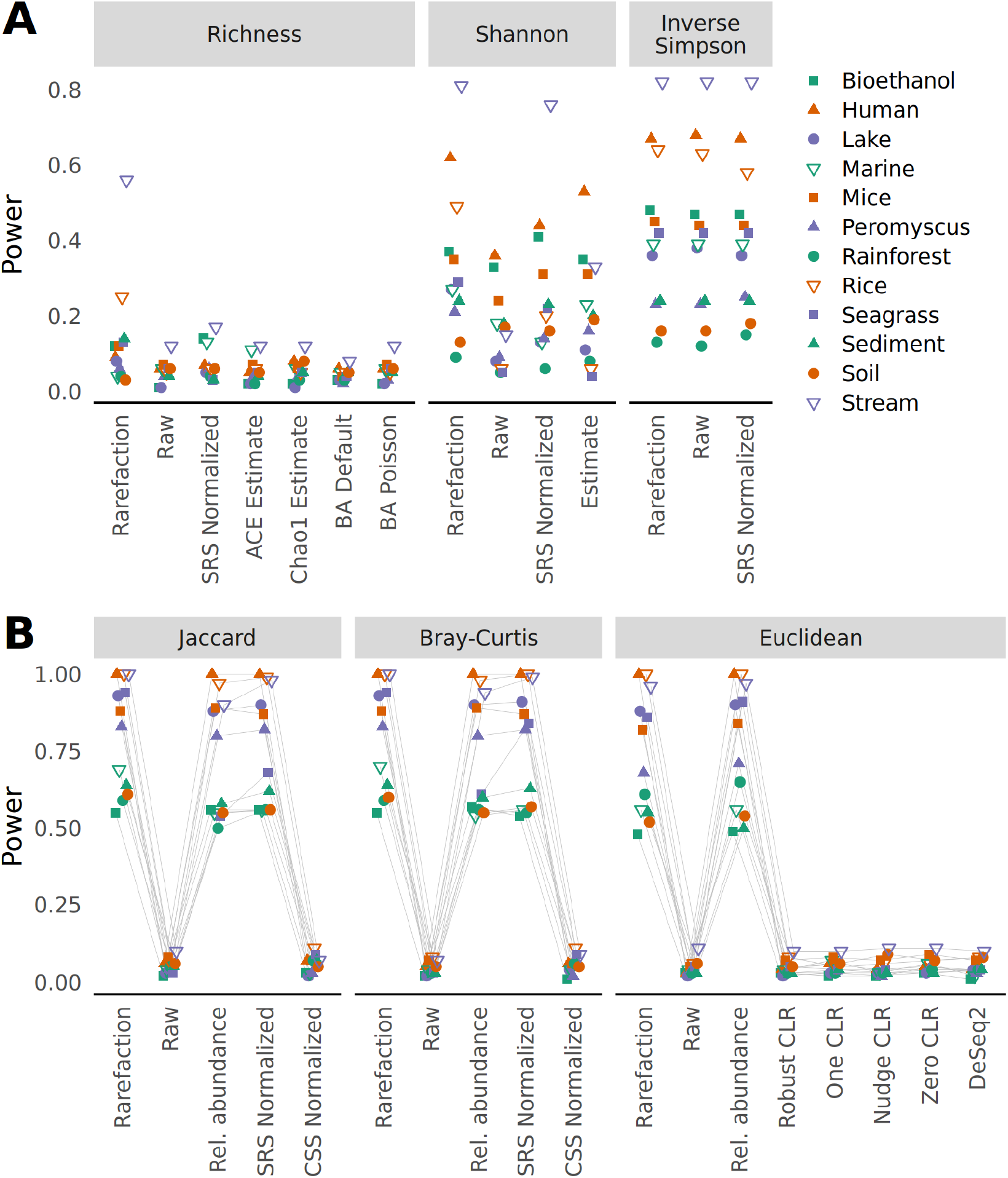
The ability to detect true differences in treatment groups for alpha (A) and beta (B) diversity metrics is greatest when communities differing in the relative abundance of their OTUs are normalized by rarefaction. For each dataset samples were randomly assigned to one of two community distributions where the abundance of OTUs differed. To calculate the power for each datasets, datasets were regenerated 100 times and differences in alpha diversity were tested using a Wilcoxon test (A) and differences in beta diversity were tested using PERMANOVA (B) at a 5% threshold. The power was the number of times a dataset yielded a significant result.

**Figure 5.**
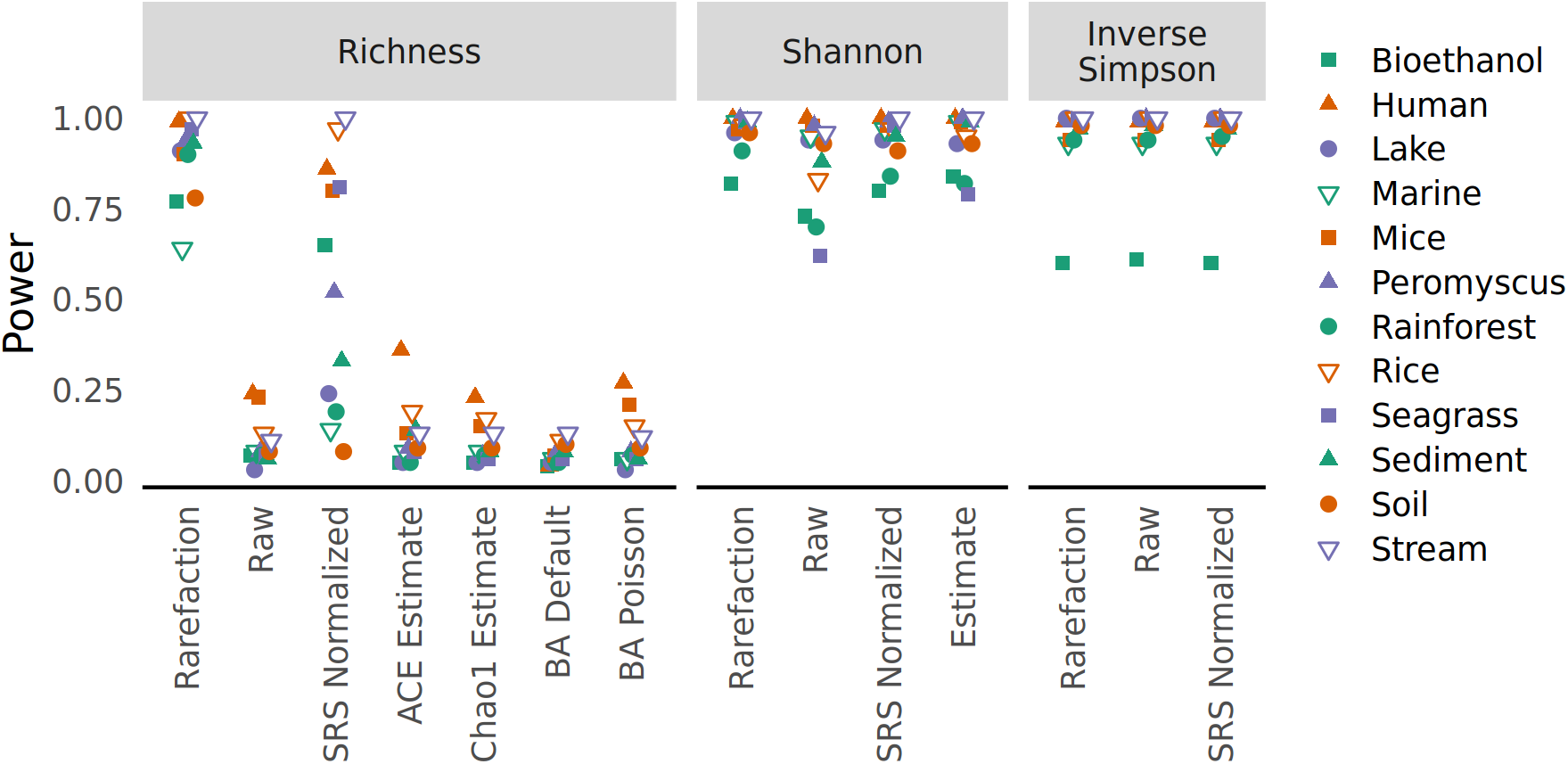
The ability to detect true differences in treatment groups for alpha diversity metrics is greatest when sequencing depths from communities differing in richness are normalized by rarefaction. For each dataset samples were randomly assigned to one of two community distributions where one distribution contained a subset of OTUs found in the other. To calculate the power for each dataset, datasets were regenerated 100 times and differences in alpha diversity were tested using a Wilcoxon test (A) and differences in beta diversity were tested using PERMANOVA (B) at a 5% threshold. The power was the number of times a dataset yielded a significant result.

### Increased rarefaction depth reduces intra-sample variation in alpha and beta diversity

Once concern with using rarefaction is the perceived loss of sequencing information when a large fraction of data appears to be removed when the community with the greatest sequencing depth is sampled to the size of the community with the least (e.g., the smallest sample in the Bioethanol dataset had 1.04% of sequences that were in the largest sample). To assess the sensitivity of alpha and beta diversity metrics to rarefaction depth, I again used the dataset generated using the null models, but used rarefaction with each community to varying sequencing depths (Figure 6). The richness values increased with sequencing effort as rare OTUs would continue be detected. In contrast, the Shannon diversity and Bray-Curtis values plateaued with increased sequencing effort. This result was expected since increased sequencing would lead to increased precision in the measured relative abundance of the OTUs. Next, I measured the coefficient of variation (i.e., the mean divided by the standard deviation) between samples for richness, Shannon diversity, and Bray-Curtis distances. Although the mean richness appeared to increased unbounded with sequencing effort, the coefficient of variation for each dataset was relatively stable. In general, the coefficient of variation increased slightly with sequencing depth only to decline once smaller samples were removed from the analysis at higher sequencing depths. Interestingly, the coefficient of variation between Shannon diversity values decreased towards zero with increased sequencing effort and the coefficient of variation between Bray-Curtis distances tended to increase. Regardless, the coefficients of variation were relatively small. This analysis indicates that there are benefits to increased sequencing depths.

**Figure 6.**
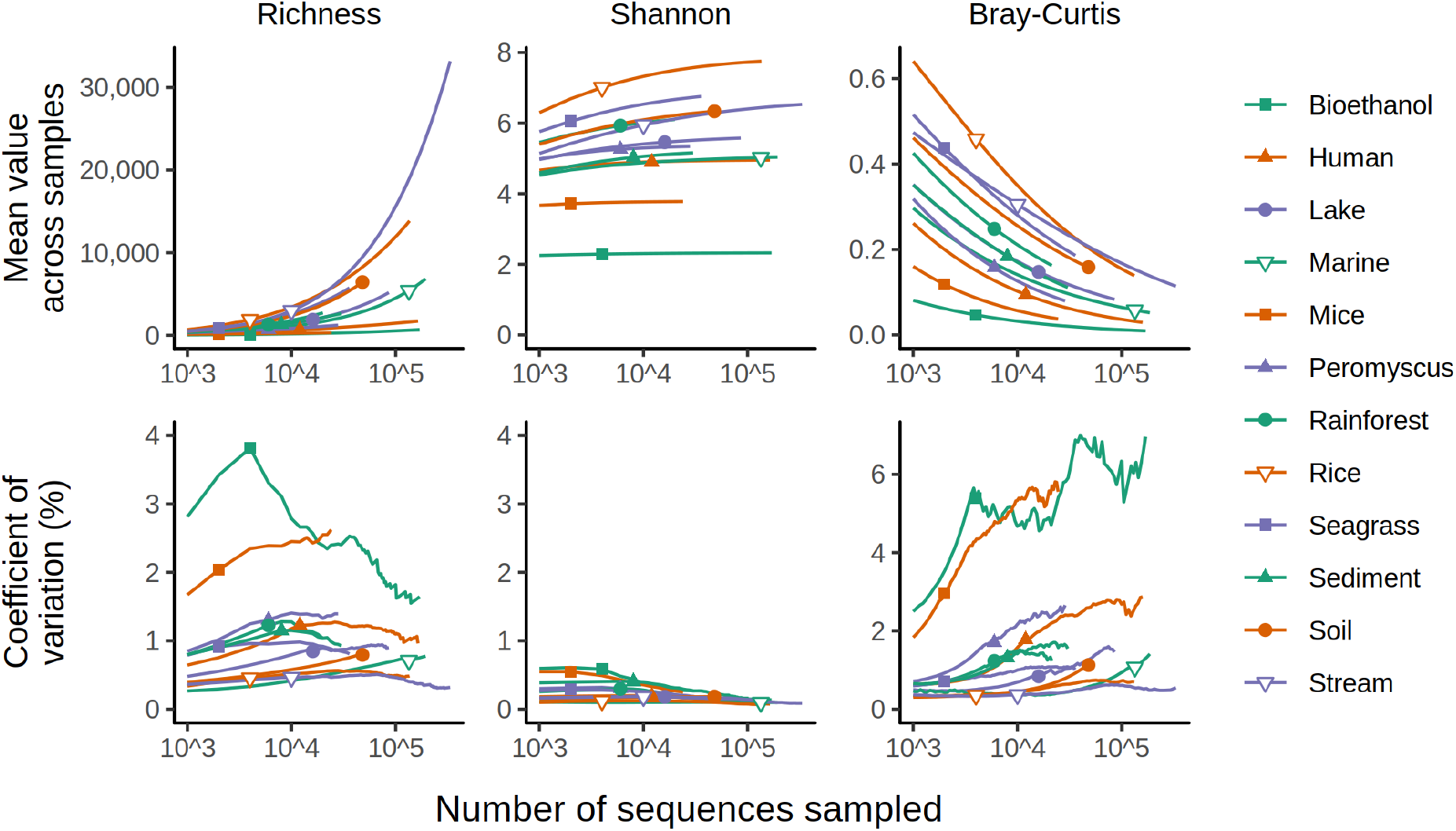
The mean and coefficient of variation for richness, Shannon diversity, and Bray-Curtis dissimilarity values calculated by rarefaction vary with sequencing depth. For each dataset, a null community distribution was created and samples were created to have the same sequencing depth as they did originally. The placement of the plotting symbol indicates the size of the smallest sample. Results are only shown for sequencing depths where a dataset had 5 or more samples.

### Most studies have a high level of sequencing coverage

To explore the concern over loss of sequencing depth further, I calculated the Good’s coverage for the observed data. The median coverage for each dataset ranged between 89.4 and 99.8% for the Seagrass and Human datasets, respectively (Figure 7). When I used a rarefaction threshold with each dataset at the size of the smallest community in the dataset, with the exception of the Seagrass, Rice, and Stream datasets, the median coverage with rarefaction was still greater than 90%. These results suggest that most studies had a level of sequencing coverage that aligned with the diversity of the communities. Next, I used the null model for each dataset to ask how much sequencing effort was required to obtain higher levels of coverage. To obtain 95 and 99% coverage, an average of 2.70 and 101.20-fold more sequence data was estimated to be required than was required to obtain 90% coverage, respectively (Figure 7). As suggested by the simulated coverages curve in Figure 7, these estimates are conservative. Regardless, the sequencing effort required to achieve higher sequencing depth would likely limit the number of samples that could be sequenced when controlling for costs. Although it may be disconcerting to use rarefaction to normalize to a sequencing depth that is considerably lower than that obtained for the best sequenced community in a dataset, sequencing coverage for many environments is probably adequate even at the lower sequencing depth. Of course, regardless of the concerns surrounding the choice of the rarefaction depth, the results throughout this study demonstrate that rarefaction is necessary to avoid reaching incorrect inferences.

**Figure 7.**
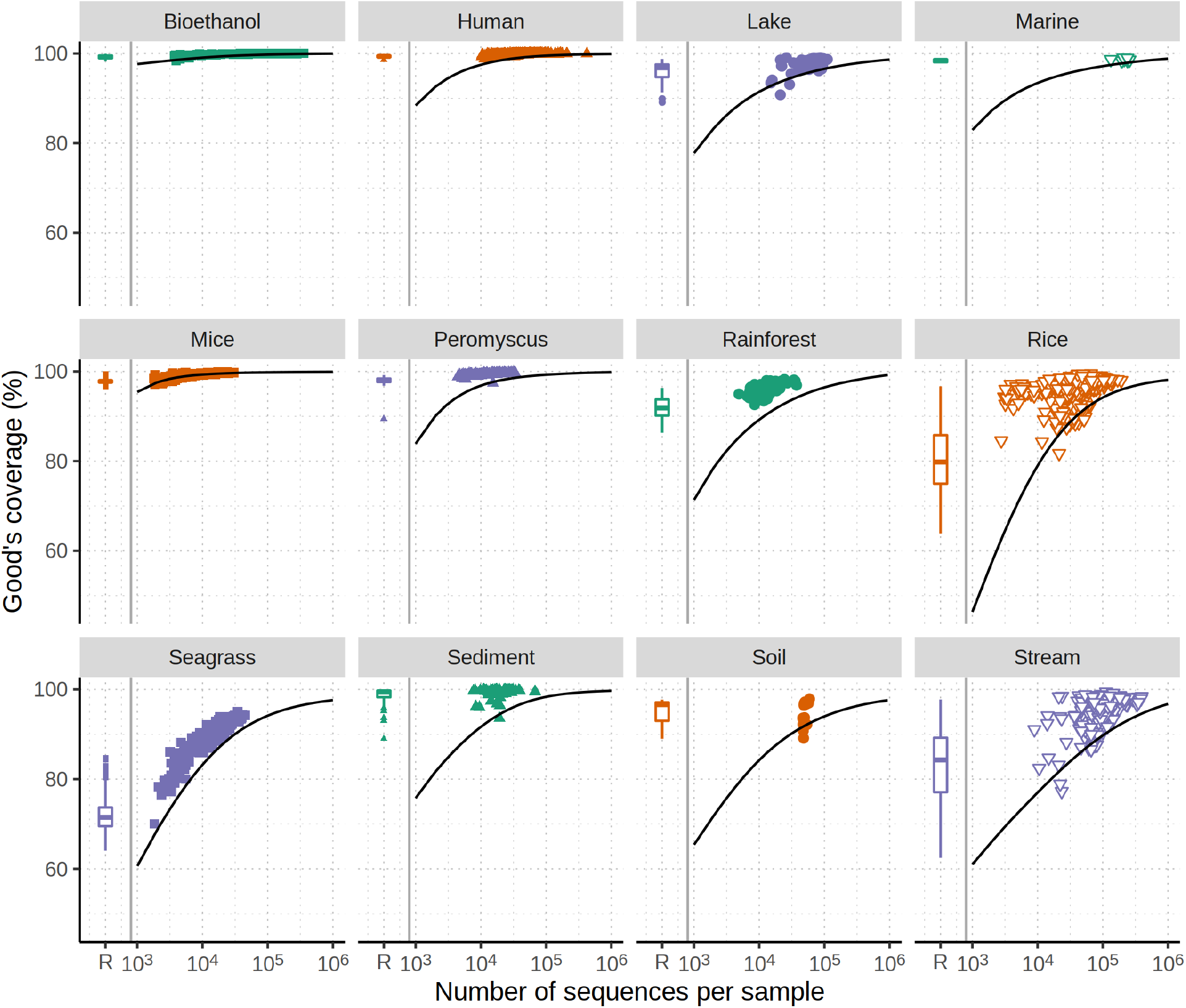
Most datasets are sequenced to a level that provides a high level of coverage. Each plotting symbol represents the observed Good’s coverage for a different sample in each dataset. The smoothed line indicates the simulated coverage for varying levels of sequencing effort when a null community is generated from the observed data. The box and whisker plot indicates the range of coverage values when the observed community data were normalized by rarefaction to the size of the least sequenced sample.

## Discussion

Over the past decade, the question of whether to use rarefaction with microbial community sequence data has become controversial. The analyses I presented here strongly indicate that rarefaction is necessary to control for uneven sequencing effort when comparing communities based on alpha and beta diversity indices. Compared to all other methods, rarefaction removed the correlation between sequencing depth and alpha or beta diversity metrics when the sequencing depth varied by as much as 97-fold across samples. I showed that this correlation could lead falsely detecting differences between treatment groups if sequencing depth and sequencing effort are confounded. The correlation with sequencing effort leads to an artificial increase in the variation between samples and a reduced power to detect true differences in alpha and beta diversity. For these reasons, rarefaction is a valuable tool to control for uneven sequencing effort until improved statistical procedures are developed or it becomes possible to more evenly distribute sequencing effort across samples.

The primary alternative to rarefaction for measuring alpha diversity is to estimate the metric using non-parametric or parametric models with raw counts and to then compare the estimates (7, 14). My results demonstrate that such approaches are limited for several reasons. First, non-parametric richness estimates such as ACE and Chao1 are sensitive to sequencing effort. Therefore, these strategies do not, in practice, remove the effects of sequencing effort. Second, parametric approaches, such as those implemented in the `breakaway` R package, generate confidence intervals that are likely to include the true richness and theoretically shrink with increased sequencing effort. Yet for most samples, the confidence intervals around the estimates are too wide to be meaningful, again leading to an inability to remove the effects of sequencing effort. Third, it has become an increasingly common practice for researchers to remove sequences that are rare in a sample (e.g., those that appear once). Although that approach was not taken in this study, removing rare sequences would skew the distribution of sequences and OTUs leading to a distortion of the measurement of alpha diversity (27). The effects of removing rare sequences would vary across samples depending on the number of sequences in each sample. One interesting result of this analysis was the demonstration that as metrics that depend less on rare taxa are used, the effect of uneven sequencing effort was reduced. For example, richness counts a sequence appearing once as much as sequence appearing 1,000 times, while the Shannon diversity index places more emphasis on the more abundant sequence, and the inverse Simpson index even more. Although normalizing communities to a common number of sequences is also suggested (e.g., SRS normalization) to control for uneven sequencing effort, the current analysis indicates that its performance does not meet that of rarefaction. For analysis of alpha diversity metrics, rarefaction is the most effective and consistent approach to control for uneven sequencing effort.

Use of relative abundances, normalized counts, variance stabilizing transformations, and centered log ratios have each been recommended as more robust alternatives to rarefaction. Again, the only approach that consistently removed the effects of uneven sequencing effort was rarefaction. Most of the recommendations borrow techniques from methods for identifying differential gene expression. Unfortunately, there is an important but under appreciated difference between gene expression and community data. This is the interpretation of unobserved data. For gene expression analysis in a single organism the lack of any sequencing data for a gene would indicate that although the gene was present, its expression was below the limit of detection. Sequencing the same organism under multiple conditions would not lead to a seemingly unbounded number of genes in the organisms. Rather, the number of genes has a definite limit that is knowable from the genome sequence. With microbiome data, an unobserved sequence could mean that the organism was present, but below the limit of detection or that the organism was missing. Because we have yet to exhaustively sample any community in the same way we have sequenced a single genome, it is unreliable to impute the presence of all organisms. Yet, this is exactly what the variance stabilization transformation and most CLR techniques do. This analysis has demonstrated a clear correlation between distances calculated by these methods and sequencing effort. This result is at odds with the claims by others that the distances are scale invariant (22, 24). Again, rarefaction is the most effective and consistent approach to control for uneven sequencing effort when calculating beta diversity metrics.

Two common critiques of rarefaction are that the approach omits valid data and that the selection of rarefaction threshold is arbitrary (11). I disagree with both critiques. All of the data are used to calculate the mean value of the metrics after repeatedly subsampling the data. When the dataset is subsampled, every sequence has a random chance of being included in the calculated metric. When that subsampling is repeated a large number of times (e.g., 100 or 1000) the risks of ignoring or oversampling rare taxa are mitigated. Furthermore, it is curious that the study making the original critique removed rare sequences. A parallel analysis to this study has demonstrated that many of these sequences are likely valid and that removal of rare sequences can bias alpha and beta diversity metrics and reduce statistical power (27). As for the second criticism, I would resist the claim that the selection of the rarefaction threshold is arbitrary. In practice, there is a tradeoff between sampling breadth and sequencing depth. Greater breadth will increase the statistical power to compare treatment groups and greater depth will increase the resolution to describe the communities. My personal process for rarefaction threshold involves looking for a natural break in the distribution of the number of sequences. For example, the Lake dataset used in this study had a clear break around 10,000 sequences (Figure S1). I would also consider what samples are below any break that I select. If there were critical samples below the break I would either reduce the threshold or obtain more sequences from those samples. As shown in my analysis of Good’s coverage values, most studies obtain an ample level of coverage and would need to increase their sequencing depth by 10-fold to increase the coverage by several percentage points. In past work, I have favored distributing the total number of sequences to increase sampling breadth rather than sequencing depth.

The up to 100-fold difference in sample sizes is an unfortunate byproduct of how sequencing libraries are constructed. Researchers perform separate PCRs for each sample with unique index (aka, barcode) sequences that allow them to later assign sequences back to the samples that they came from. When the PCR products are pooled, efforts are taken to pool the fragments in equimolar ratios. Researchers use one of two approaches. First, they often will quantify the concentration of DNA from each PCR and then pool DNA in the desired amounts. Alternatively, they may use normalization plates where each well can hybridize a uniform amount of DNA that is then eluted and pooled. Clearly, both approaches have limitations that reduce the ability to truly achieve equimolar mixture. In addition, for some samples it is common to co-amplify non-specific DNAs which may add to the challenges of achieving equimolar mixtures of the desired amplicons since those sequences will be removed (28). Regardless, it is clear that better strategies are needed to reduce the variation in the number of sequences generated for each sample.

All simulations have weaknesses and should be interpreted with caution. However, the simulated communities generated and analyzed in this study had the advantage of being designed with known properties including the alpha and beta diversity and the their differences between treatment groups. Indeed, is is perfectly admissible and proper to use rarefaction with microbial community data. The alternative is to risk reaching unwarranted conclusions.

## Materials and Methods

### Choice of datasets

The specific studies used in this study were selected because their data was publicly accessible through the Sequence Read Archive, the original investigators sequenced the V4 region of the 16S rRNA gene using paired 250 nt reads, and my previous familiarity with the data. The use of paired 250 nt reads to sequence the V4 region resulted in a near complete two-fold overlap of the V4 region resulting in high quality contigs with a low sequencing error rate (3). These data were processed through a standardized sequence curation pipeline to generate operational taxonomic units (OTUs) using the mothur software package (3, 9). OTUs were identified using the OptiClust algorithm to cluster sequences together that were not more than 3% different from each other (29).

### Null community model

Null community models were generated such that within a dataset the number of sequences per sample and the number of sequences per OTU across all samples within the dataset were the same as was observed in original. This model effectively generated statistical samples of a single community so that there should not have been a statistical difference between the samples. This model implemented by randomly assigning each sequence in the dataset to an OTU and sample while keeping constant the number of sequences per sample and the total number of sequences in each OTU. This is a similar approach to that of the IS algorithm described by Ulrich and Gotelli (30). Because the construction of the null models was a stochastic process, 100 replicates were generated for each dataset.

### Null treatment models

I created an unbiased and biased treatment model. In the unbiased model, samples were randomly assigned to one of two treatment groups. In the biased treatment model, samples that had more than the median number of sequences for a dataset were assigned to one treatment group and the rest were assigned to a second treatment group. Comparison of any diversity metric across the two treatment groups should have only yielded a significant result in 5% of the simulations when testing under a Type I error (i.e., *α*) of 0.05.

### Skewed abundance community model

In the skewed abundance community model, communities were randomly assigned to one of two simulated treatment groups. Communities in the first treatment group were generated by calculating the relative abundance of each OTU across all samples and using those values as the probability of sampling each OTU. This probability distribution was sampled, with replacement, until each sample had the same number of sequences that it did in the observed data. Samples in the second treatment group were generated by adjusting the relative abundances of the OTUs in the firs treatment group by increasing the relative abundance of 10% of the OTUs by 5%. These values were determined after empirically searching for conditions that resulted in a large fraction of the randomizations yielding a significant result across most of the studies. Sequences were sampled from the skewed community community until each sample had the same number of sequences that it did in the observed data. Under the skewed abundance community model, each sample represented a statistical sampling of two communities such that there should not have been a statistically significant difference within a treatment group, but there was between the treatment groups. Because the construction of the skewed abundance community model was a stochastic process, 100 replicates were generated for each dataset.

### Richness-adjusted community model

In the richness-adjusted community model, communities were randomly assigned to one of two simulated treatment groups. Communities in the first treatment group were generated by calculating the relative abundance of each OTU across all samples and using those values as the probability of sampling each OTU. This probability distribution was sampled until each sample had the same number of sequences that it did in the observed data. Samples in the second treatment group were generated by removing 3% of the OTUs from the dataset and recalculating the relative abundance of the remaining OTUs. Sequences were sampled from the richness-adjusted community distribtion, with replacement, until each sample had the same number of sequences that it did in the observed data. Under the richness-adjusted community model, each sample represented a statistical sampling of two communities such that there should not have been a statistically significant difference within a treatment group, but there was between the treatment groups. Because the construction of the richness-adjusted community model was a stochastic process, 100 replicates were generated for each dataset.

### Test of statistical significance

Statistical comparisons of alpha diversity metrics across the simulated treatment groups were performed using the non-parametric two-sample Wilcoxon test as implemented in `wilcoxon.test` in the `stats` base R package. This test was selected because the alpha-diversity metrics tended to not be normally distributed and each dataset required a different transformation to normalize the data. Comparisons of beta diversity metrics were performed using the `adonis2` function from the `vegan` (v.2.6.2) R package (10). The `adonis2` function implements a non-parametric multivariate analysis of variance using distances matrices (31). Throughout this study I used 0.05 as the threshold for assessing the statistical significance of any P-values.

### Power analysis

The parameters used to design the skewed abundance and richness-adjusted community models were set to impose a known effect size when using community data normalized by rarefaction. The statistical power to detect these differences was determined by calculating the p-value for each of 100 replicate simulated set of samples from each dataset using the various alpha and beta diversity metrics. The percentage of tests that yielded a significant P-value was considered the statistical power (i.e., 1 minus the Type II error) to detect the difference.

### Alpha diversity calculations

Various strategies for handling uneven sequencing effort were evaluated to identify the best approach for calculating community richness and Shannon and inverse Simpson diversity indices. OTU counts were used as input to calculate sample richness and Shannon and inverse Simpson diversity using mothur (9, 32). Shannon diversity was calculated as

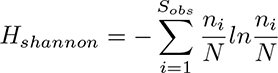

The Simpson diversity was calculated as

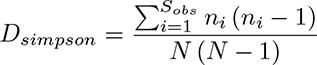

The inverse Simpson diversity was calculated as `1`*/D_simpson_*. In both formulae, *n_i_*was the number of sequences in OTU *i* and *N* iwass the number of sequences in the sample. Rarefaction of richness, Shannon diversity and Inverse Simpson diversity values were carried out in mothur. Briefly, mothur calculates each value on a random draw of the same number of sequences from each sample and obtains a mean value based on 1,000 random draws. Scaled ranked subsampling (SRS) was used to normalize OTU counts to the size of the smallest sample in each dataset using the SRS R package (v.0.2.3)(19). Normalized OTU counts were used to calculate sample richness and Shannon and inverse Simpson diversity values using mothur. Data normalized by cumulative sum scaling (CSS) were not reported for alpha-diversity values since the relative abundances of the features do not change with the normalization procedure (20). The non-parametric bias-corrected Chao1 and ACE richness estimators (16) and a non-parametric estimator of the Shannon diversity (15) were calculated using raw OTU counts with mothur. Parametric estimates of sample richness were calculated using the `breakaway` (BA) R package (v.4.7.9)(17). My analysis reports both the results from running default model selection procedure and the Poisson model. The default model selection returned either the Kemp, Negative Binomial, or Poisson models. Relative abundance data were not used to calculate alpha diversity metrics since the richness and evenness does not change from the raw data when dividing each sample by the total number of sequences in the sample.

### Beta diversity calculations

Similar to the alpha diversity calculations, multiple approaches were used to control for uneven sequencing effort and calculate beta diversity. Raw and OTU counts were used for input to calculate the Jaccard, Bray-Curtis, and Euclidean dissimilarity indices using the `vegdist` function from the `vegan` R package (v.2.6.2)(10). The Jaccard index was calculated as

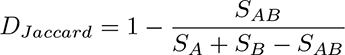

where *S_A_*and *S_B_*were the number of OTUs in samples *A* and *B* and *S_AB_*was the number of OTUs shared between the two samples. The Bray-Curtis index was calculated as

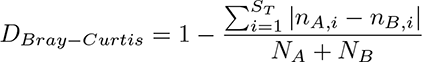

where *n_A,i_*and *n_B,i_*were the number of sequences observed in OTU *i* from samples *A* and *B*, respectively. *N_A_* and *N_B_* were the total number of sequences in samples *A* and *B*, respectively. *S_T_* was the total number of OTUs observed between the two samples. The Euclidean distance was calculated as

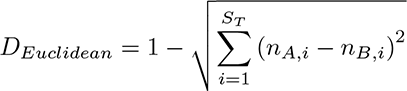

These metrics were calculated using the relative abundance of each OTU using the `vegdist` function from the `vegan` R package. The relative abundance was calculated as the number of sequences in the OTU (e.g., *n_A,i_*) divided by the total number of sequences in the sample (e.g., *N_A_*).

Beta-diversity values generated with rarefaction were calculated using the `avgdist` function in vegan. Briefly, `vegan`’s `avgdist` function calculates each pairwise dissimilarity index after obtaining a random draw of the same number of sequences from each sample. After obtaining 100 random draws it returns the mean value.

Three approaches were taken to normalize the number of sequences across samples within a dataset. Scaled ranked subsampling (SRS) and cumulative sum scaling (CSS) were used to normalize raw OTU counts using the `SRS` (v.0.2.3) and `metagenomeSeq` (v.1.36.0) R packages, respectively (19, 20). The normalized counts were then used to calculate Jaccard and Bray-Curtis dissimilarity indices. Finally, the variance-stabilization transformation (VST) as implemented in the `DESeq2` (v.1.34.0) R package was used to normalize the data as described by McMurdie and Holmes (11, 33). Because the VST approach generated negative values, which are incompatible with calculating Jaccard and Bray-Curtis dissimilarity values, Euclidean distances were calculated instead.

Raw OTU counts were used to calculate centered log ratio (CLR) values for each OTU, which were then used to calculate Euclidean distances; such distances are commonly referred to as Aitchison distances. CLR abundances are calculated as:

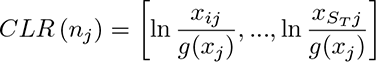

where *x_ij_* was the number of sequences observed for OTU *i* in sample *j* and *g*() was the geometric mean, *x_j_*was the count of the *S_T_* OTUs in sample *j*. Because the geometric mean is zero if any OTU is absent from a sample, the CLR is undefined when there are unobserved OTUs in a sample. To overcome this problem, I attempted a four approaches. The first, Zero CLR, imputed the value of the zeroes based on the observed data using the zCompositions (v.1.4.0.1) R package (34). The second, One CLR, added a pseudo-count of 1 to the abundance of all OTUs (18, 20). The third, Nudge CLR, added a pseudo-count of 1 divided by the total number of sequences in a sample to each OTU in the sample (18, 23). The final approach, Robust CLR, removed unobserved OTUs prior to calculating the CLR (22).

### Analysis of sequencing coverage

To assess the level of sequencing coverage I calculated Good’s coverage (*C_Good_*) using mothur:

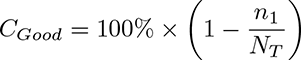

where *n*_1_ was the number of OTUs with only one sequence in the sample and *N_T_* was the total number of sequences in the sample. Good’s coverage was calculated using (i) the observed OTU counts for each sample and dataset, (ii) following rarefaction (1,000 iterations) of the observed OTU counts to the size of the smallest sample in each dataset, and (iii) after rarefaction (1,000 iterations) of the null community distribution.

### Reproducible data analysis

A complete reproducible workflow written in Snakemake (v.7.15.2) and Conda (v.4.12.0) computational environment can be obtained from the GitHub hosted git repository for this project (https://github.com/SchlossLab/Schloss_Rarefaction_XXXXX_2023). This paper was written in R-flavored markdown (v.2.16) with the `kableExtra` (v.1.3.4) package. The mothur (v.1.47.0) and R (4.1.3) software packages were used for all analyses with extensive use of functions in the tidyverse metapackage (v.1.3.1). A preliminary version of this analysis was presented as the Rarefaction video series on the Riffomonas Project YouTube channel (https://www.youtube.com/playlist?list=PLmNrK_nkqBpJuhS93PYC-Xr5oqur7IIWf).

## Acknowledgements

I am grateful to the researchers who generated the datasets used in this study. I also thank the individuals who asked questions and commented on the preliminary version of this project, which was released as a YouTube playlist on the Riffomonas channel. This work was supported in part by funding from the National Institutes of Health (U01AI124255, P30DK034933, R01CA215574).

## Figures

**Figure S1.**
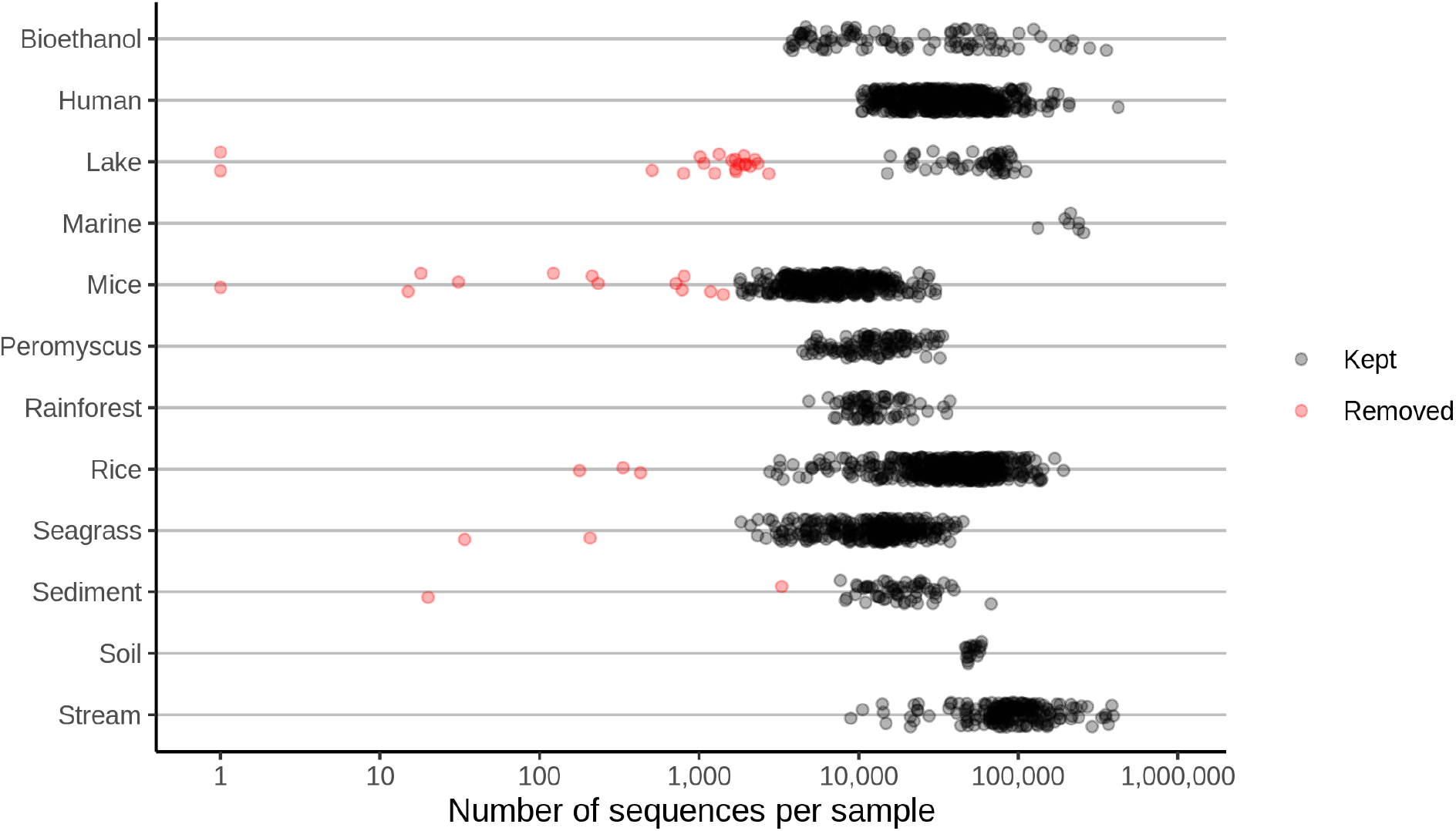
The number of sequences observed in each sample for each dataset included in this analysis generally varied by 10 to 100-fold. The threshold for specifying the number of sequences per sample varied by dataset and was determined based on identifying natural breaks in the data. This figure is similar to Figure S1 of (27)

**Figure S2.**
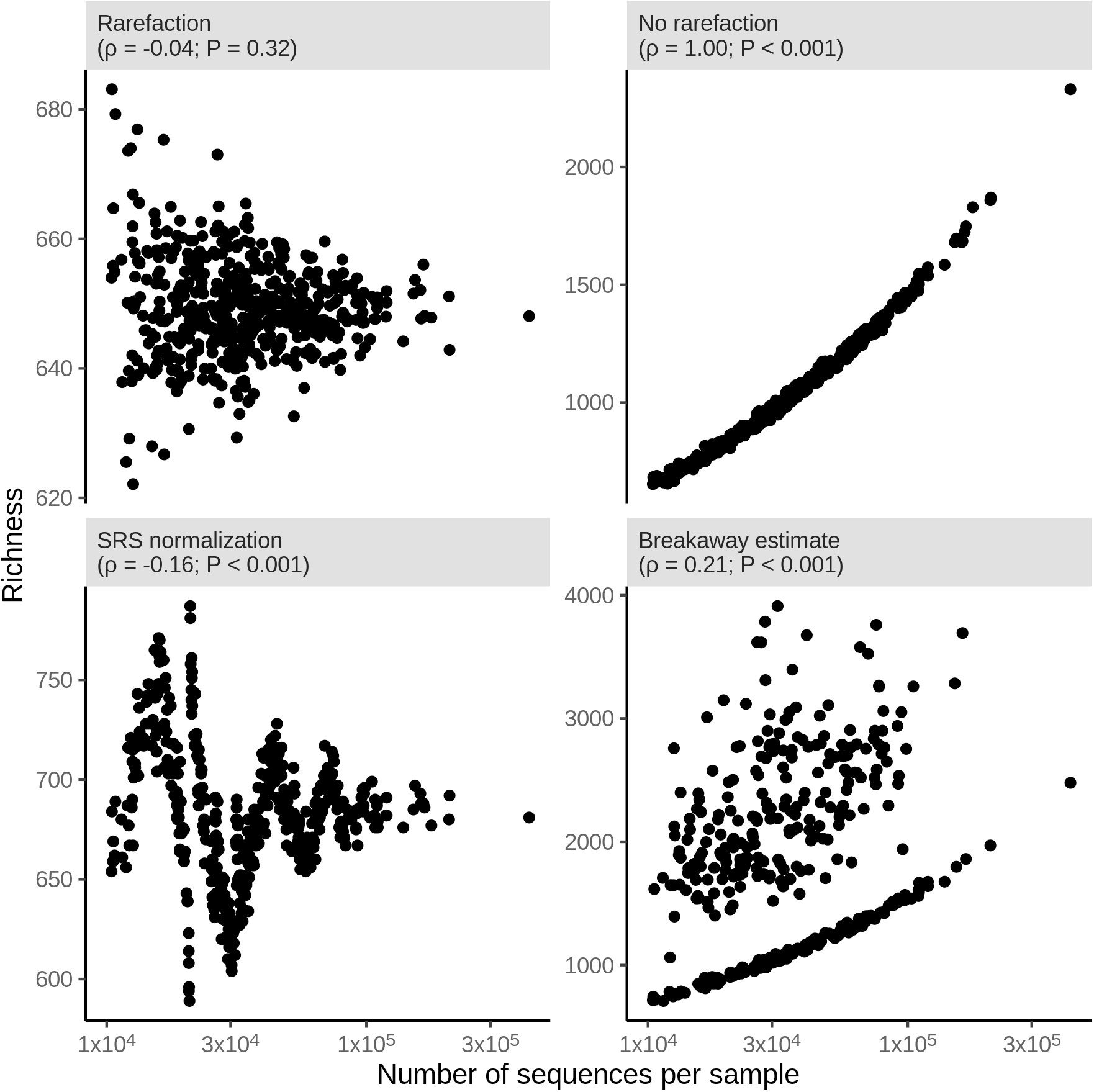
Examples of the richness in each of the 490 samples that were generated for one randomization of the null model using the human dataset. The x-axis indicates the number of sequences in each of the samples prior to each method’s approach of controlling for uneven sequencing effort. The Spearman correlation coefficient (*ρ*) and test of whether the coefficient was significantly different from zero are indicated for each panel.

**Figure S3.**
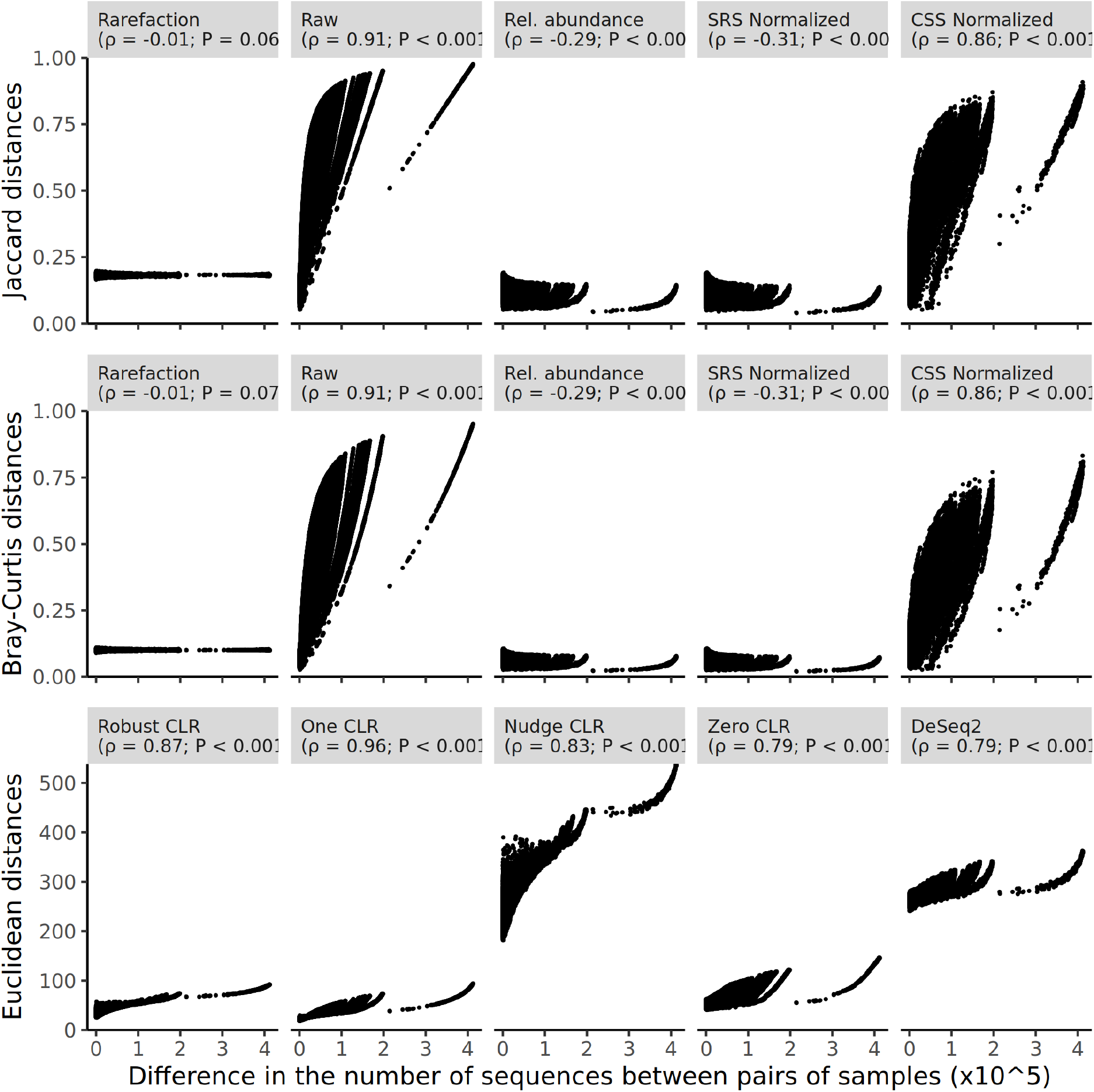
Examples of differences in beta diversity in each of the 490 samples that were generated for one randomization of the null model using the human dataset. The x-axis indicates the difference in the number of sequences in each of the samples that went into calculating the pairwise distance prior to each method’s approach of controlling for uneven sequencing effort. The Spearman correlation coefficient (*ρ*) and test of whether the coefficient was significantly different from zero are indicated for each panel.

## Notes

### Competing Interest Statement

The authors have declared no competing interest.

